# OligoGraph: A novel geometric graph-based approach for siRNA efficacy prediction

**DOI:** 10.64898/2026.02.23.707354

**Authors:** Sriram Shravan Saligram, Vishnu Vardhan Kasturi, Sidhardha Reddy Surkanti, Bhargava Chary Basangari, Vani Kondaparthi

**Author notes:** Correspondence: Vani Kondaparthi Correspondence email ID. Authors contributed equally (Sriram Shravan Saligram, Vishnu Vardhan Kasturi, Sidhardha Reddy Surkanti).

## Abstract

RNA interference (RNAi) is a biological process in which a small interfering RNA (siRNA) prevents the translation of a messenger RNA (mRNA) into a protein by cleaving the mRNA before translation. We exploit this process to prevent the formation of harmful proteins by using an effective siRNA on the target mRNA. The current rapidly emerging RNAi-based drugs show immense potential for therapeutic applications.

Traditionally, designing a potent siRNA for an mRNA requires extensive lab experimentation and trials; therefore, there is a need to develop a model that reliably predicts a siRNA’s efficacy against mRNA. This saves both cost and time. But designing such models is challenging, as the data available is either scarce or biased. The current models available exhibit limited generalization and are restricted to a fixed siRNA lengths of either 19 or 21 nucleotides, limiting flexible use.

To address these challenges, we introduce OligoGraph, a graph-based deep learning architecture that operates on the siRNA-mRNA duplex. It leverages RiNALMo embeddings, multiple GATconv and Transformerconv layers, and self-supervised pretraining, and outperforms all other existing models in our testing on seen and unseen data.

We implemented specialized OligoGraph variants for 19- and 21-nucleotide siRNAs, both of which outperformed the current state-of-the-art models on unseen data. The 19-nucleotide model yielded AUC-ROC and PCC increases of 1.1% and 4.6% on the Mixset; 19.07% and 127.3% on the Takayuki dataset, respectively. Furthermore, the 21-nucleotide model improved predictive performance on the Simone dataset by 2.62% (AUC-ROC) and 6.65% (PCC).

## 1. Introduction

RNA interference (RNAi) is a natural biological gene-silencing mechanism in which siRNA molecules regulate gene expression by degrading sequence-specific mRNA, thereby preventing the respective protein formation.

The process starts when a double-stranded RNA (dsRNA) enters the cytoplasm and gets processed by dicer, an RNase III endonuclease, which slices the dsRNA into two 21-25 nucleotide siRNAs containing 2-nucleotide 3’ overhangs. These siRNAs interact with the RNA-induced silencing complex, which contains Argonaute proteins, especially Ago2. During RISC loading thermodynamic asymmetry decides the guide strand selection with the strand showing reduced 5’ stability preferably loaded as the guide strand while the passenger strand gets degraded [1]. The RISC, guided by the guide strand, then targets complementary mRNA through Watson-Crick pairing [2], leading to Ago2-mediated mRNA cleavage [3]. This cleavage results in rapid exonucleolytic degradation of the mRNA fragments [4], thereby removing the template for protein synthesis and silencing gene expression at the post-transcriptional level.

Optimal siRNA design requires thermodynamic features to ensure correct anti-sense strand loading with the 5’ end of the antisense strand showing lower stability than the sense 3’ end. Currently 8 RNAi (siRNA) drugs have been approved by FDA and with over 260 drug candidates in development as of 2025 (https://www.cas.org/resources/cas-insights/sirna-therapeutics).

Critical challenges include off-target effects by siRNAs interacting with unintended mRNA targets, activation of immune responses, and delivery barriers. Additionally, siRNA design requires extensive computational design and validation to optimize guide strand selection and minimize passenger strand retention. Hence, predicting effective siRNAs is crucial for RNAi therapeutics, leading to the development of computational tools to aid this process.

The first generation of siRNA computational tools, like Reynold’s algorithm [5], Ui-Tei rules [6], Amarzguioui guidelines [7] relied on observations from small, validated datasets to create position-specific rules for siRNA efficacy prediction.

The second-generation models transitioned from rule-based algorithms to machine learning models. BIOPREDsi [8] in 2005, developed by Huesken et al., used a Stuttgart neural net simulator to train artificial neural networks. I-SCORE [9] in 2007 employed linear regression along with thermodynamic instability requirements. OligoWalk [10] in 2008 used partition function calculations with hybridization thermodynamics for their tool. DSIR [11] in 2012 used linear regression models with position-specific nucleotide preferences for siRNA efficacy prediction. VIRsiRNApred [12] in 2013 used support vector machines along with thermodynamic properties and nucleotide frequencies to target 1725 experimentally verified viral siRNAs against 37 human viruses to train their model. SMEpred [13] in 2016 was developed to predict the efficacy of normal siRNAs along with chemically modified siRNAs using support vector machines.

The third generation models transitioned from machine learning models to deep learning architectures. HAN [14] in 2017 employed CNN predictions. GNN4siRNA [15] in 2024 introduced the first ever graph neural network approach for siRNA efficacy prediction to capture mRNA and siRNA molecule interactions. siRNADiscovery [16] in 2024 used a comprehensive GNN architecture integrating empirical and non-empirical features

The fourth-generation models transitioned to transformer-based models. OligoFormer [17] in 2024 used the transformer architecture with three modules thermodynamic calculations, RNA-FM a pretrained transformer encoder only module and OligoEncoder which consists of CNN, BiLSTM and transformer encoder layers to achieve current state of the art performance in inter and intra dataset metrics comparison.

Due to the scarcity of open-source RNAi datasets, there exists a need to integrate pretrained RNA foundation models to improve model generalization across datasets.

RNA-FM [18] in 2022 used the transformer architecture and was trained on 23+ million non-coding (ncRNA) sequences through self-supervised learning.

RiNALMo [19] employed modern architectural improvements of the transformer in 2025. It was trained on 36 million non-coding RNA sequences (ncRNA) from RNAcentral representing the best pre-trained model for its generalization capabilities for downstream tasks.

Here, we propose OligoGraph, a novel method for siRNA efficacy prediction. It leverages pretrained RiNALMo embeddings to capture rich nucleotide level representations enabling superior generalization across diverse siRNA – mRNA sequence contexts. The architecture employs multiple graph convolutional layers including GATConv and TransformerConv, each utilizing distinct attention mechanisms to model the complex spatial and structural relationships within the siRNA – mRNA pair. By integrating carefully selected physiochemical siRNA features such as thermodynamic stability, base pairing properties and sequence composition, OligoGraph comprehensively captures the interactions that effect the RNAi efficacy. This hybrid approach allows the model to effectively predict siRNA silencing potential by combing biological domain knowledge with data driven feature learning, providing accurate efficacy prediction.

## 2. Methods

### 2.1. Data collection and preprocessing

#### 2.1.1. Dataset collection

A diverse array of publicly available and validated datasets was collected for accuracy and robustness of our model. 9 datasets were utilized containing 3,714 siRNAs of 19 and 21 nucleotides and 75 mRNAs from previous studies such as Huesken [8], Takayuki [20], Amarzguioui [21], Harborth [22], Hsieh [23], Khvorova [1], Reynolds [5], Vickers [24], and Ui-Tei [6]. The datasets for 19-nucleotide version model were then divided into three sets as presented in Table 1, the Huesken dataset, the Takayuki dataset, and merged the remaining datasets into a single set, the Mixset. The efficacy values of siRNAs across all datasets were normalized into efficacy ranging from 0% to 100%, and 70% of the original efficacy was used as a threshold to classify siRNAs as effective, and ineffective siRNAs.

**Table 1.**
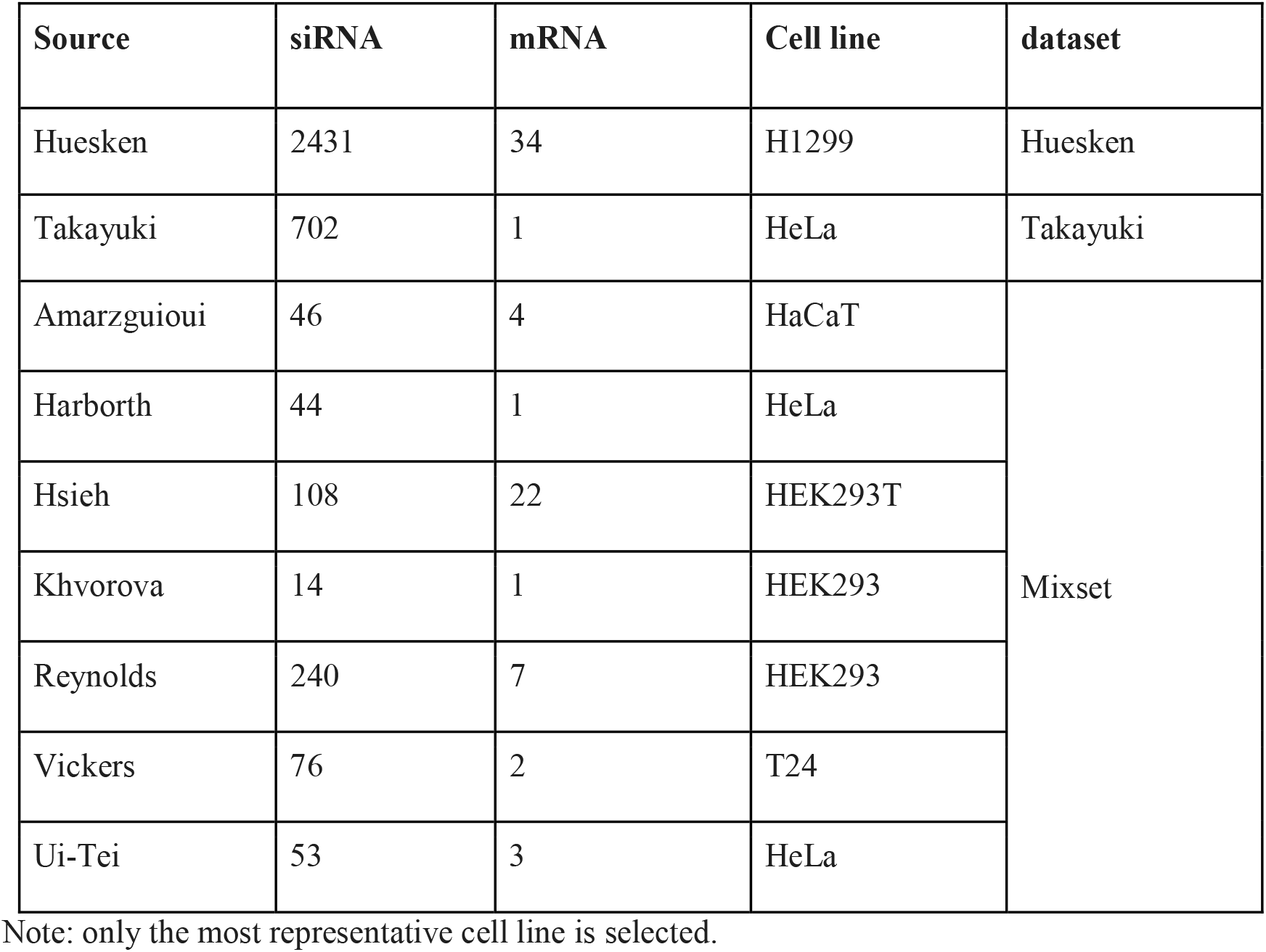
siRNA datasets.

| Source | siRNA | mRNA | Cell line | dataset |
| --- | --- | --- | --- | --- |
| Huesken | 2431 | 34 | H1299 | Huesken |
| Takayuki | 702 | 1 | HeLa | Takayuki |
| Amarzguioui | 46 | 4 | HaCaT | Mixset |
| Harborth | 44 | 1 | HeLa |  |
| Hsieh | 108 | 22 | HEK293T |  |
| Khvorova | 14 | 1 | HEK293 |  |
| Reynolds | 240 | 7 | HEK293 |  |
| Vickers | 76 | 2 | T24 |  |
| Ui-Tei | 53 | 3 | HeLa |  |

Note: only the most representative cell line is selected.

##### The Huesken Training Dataset

The Huesken training dataset contains 2,361 siRNAs generated by Huesken et al. using standardized high-throughput dual-luciferase reporter assays. All experiments were conducted on H1299 human non-small cell lung carcinoma cells at a consistent 100 nM siRNA concentration. This dataset shows a nearly normal efficacy distribution with a mean knockdown efficiency of 0.51(standard deviation), minimal skewness of −0.064, and mean GC content of 50.37%.

##### The Mixset Testing Dataset

In contrast, the Mixset testing dataset aggregates 472 siRNAs from multiple independent studies. These experiments used diverse cell lines (HeLa, HaCaT, HEK293, HEK293T, and T24) and employed various methodological approaches, including quantitative RT-PCR, Western blot protein quantification, and different reporter gene assays with varying normalization protocols. The siRNA concentrations also differed substantially across studies: Reynolds used 100 nM, Harborth used 10 nM, and Hsieh and Vickers used 20-50 nM. This heterogeneous dataset exhibits a different efficacy distribution with a mean of 0.69 and a negative skewness of −0.770, indicating enrichment toward highly effective siRNAs. The mean GC content is 49.48%.

##### Statistical Differences between Datasets

Statistical analyses reveal significant divergence between the two datasets across multiple dimensions. Independent t-tests show dramatically different efficacy means (p=5.67×10−78), while Levene’s test confirms unequal variances (p=4.02×10−69). The Kolmogorov-Smirnov test demonstrates different distribution shapes (p=2.98×10−90), and GC content distributions also differ significantly (p=4.55×10−2).

The concentration differences warrant particular attention. Higher concentrations like 100 nM can mask sequence-specific effectiveness through saturation effects and off-target modulation. Lower concentrations between 3-30 nM provide better discriminatory power for distinguishing highly functional siRNAs from moderately functional ones, since efficacy measurements follow dose-response relationships.

By training the model on the homogeneous Huesken dataset and testing it on the heterogeneous Mixset, we can evaluate how well the model generalizes across real-world variations. These include cell-line-specific biological differences, diverse experimental methodologies, and concentration-dependent effects that researchers encounter when working with siRNA efficacy data.

For the 21-nucleootide version model the datasets used were Huesken, Harborth, Ui-Tei, Vickers and Khvorova. We named this dataset HUVK. Additionally, we used the Simone dataset [25] to further validate our model.

| Dataset | siRNA | mRNA |
| --- | --- | --- |
| HUVK | 2816 | 44 |
| Simone | 322 | 6 |

#### 2.1.2. Sequence normalization

The datasets collected differ in length for siRNA and mRNA. For model training, it is crucial to ensure that siRNAs and mRNAs across the datasets have consistent lengths. The siRNA lengths in some datasets were 19 base pairing nucleotides with 2-nucleotide 3’ overhangs, while the rest contained only 19 base pairing nucleotides. To ensure consistent siRNA length for model training, we trained two versions of the model, one with 19-nucleotide sequences and the other with 21-nucleotide sequences. For the 19-nucleotide version, we have truncated the siRNA sequences to 19 nucleotides by removing 2-nucleotide 3’ overhangs. For the 21-nucleotide version, datasets with only 19-nucleotide siRNA length were excluded.

The mRNA sequence was also truncated, retaining only the binding site of the siRNA and removing the remaining nucleotides to maintain consistent mRNA length.

#### 2.1.3. RiNALMo model embeddings

In this study, The RNA language model, RiNALMo is used to generate embeddings for the siRNA-mRNA duplex [19]. The model generates 1280-dimension nucleotide level embeddings for both siRNA and mRNA. RiNALMo aids OligoGraph by capturing structural and evolutionary information at a nucleotide level. The RiNALMo model was trained on 36 million RNA sequences which helps enhance OligoGraph’s generalization capabilities thereby outperforming other models on unseen data. The model’s transfer learning foundation reduces the need for large, labeled siRNA datasets while maintaining superior generalization.

#### 2.1.4. siRNA features

The model integrates thirty thermodynamic, structural and positional features from siRNA sequences providing valuable insights into the thermodynamic stability and binding affinity of siRNA-mRNA and their interactions to enhance siRNA efficacy prediction. The features include position-specific dinucleotide Gibbs free energy (ΔG) calculations at terminal positions derived using SantaLucia/Turner [26-28] nearest-neighbor parameters, capturing energy patterns critical for binding efficacy. ΔH calculations for heat absorptions during duplex formation. A 5’ and 3’ end ΔG difference score, as the thermodynamic asymmetry affects which siRNA strand is selected as the guide strand or the passenger strand. A melting temperature (Tm) and global GC content along with seed region GC content are calculated. Five positional nucleotide identity flags and five positional dinucleotide identity flags are calculated along with global composition scores.

#### 2.1.5. Graph construction

For the model, each siRNA-mRNA interaction is constructed into a graph G = (V, E) such that, Nodes (V): Nucleotides from the siRNA guide strand and nucleotides from the mRNA target strand are represented as nodes in the graph.

Edges (E): The edges are designed to represent two types of relationships in the duplex:

1. Intra-strand edges or backbone edges: Connecting consecutive nucleotides within each strand via bidirectional connections (i, i+1) and (i+1, i) for i ∈ [0, N−2] (phosphodiester bonds) [29].
2. Inter-strand edges or base-pairing edges: Connects complementary positions between siRNA and mRNA strands. For each position i ∈ [0, min(*N*_siRNA_,*N*_mRNA_)], bidirectional edges connect siRNA node i to mRNA node i. This models the Watson-Crick and wobble base pairing across the siRNA-mRNA duplex.

Edge features: Edge features include backbone indicators, base-pairing indicators, one-hot encoding of pairing types, canonical pairing flag, wobble pairing flag, thermodynamic stability score and a seed region indicator.

Node features: Each node is the concatenation of a one-hot encode projection and RiNALMo pre-trained RNA language model embedding projection.

### 2.2. Model architecture

#### 2.2.1. Input feature processing

The RiNALMo model embeddings (*X*_*RiNALMo*_ ∈ *R*^*N*×1280^) and one-hot encodings (*X*_*Onehot*_ ∈ *R*^*N*×4^) are calculated from siRNA and mRNA, which are used as input features for the model as presented in the model overview Figure 1. Projections of these features are calculated using linear layers with layer normalization and GELU activation to get RiNALMo embedding projection (*H*_*RiNALMo*_) and one-hot embedding projection (*H*_*Onehot*_). The projections are then concatenated to create a single node-wise embedding (*H*_*node*_).

**Figure 1.**
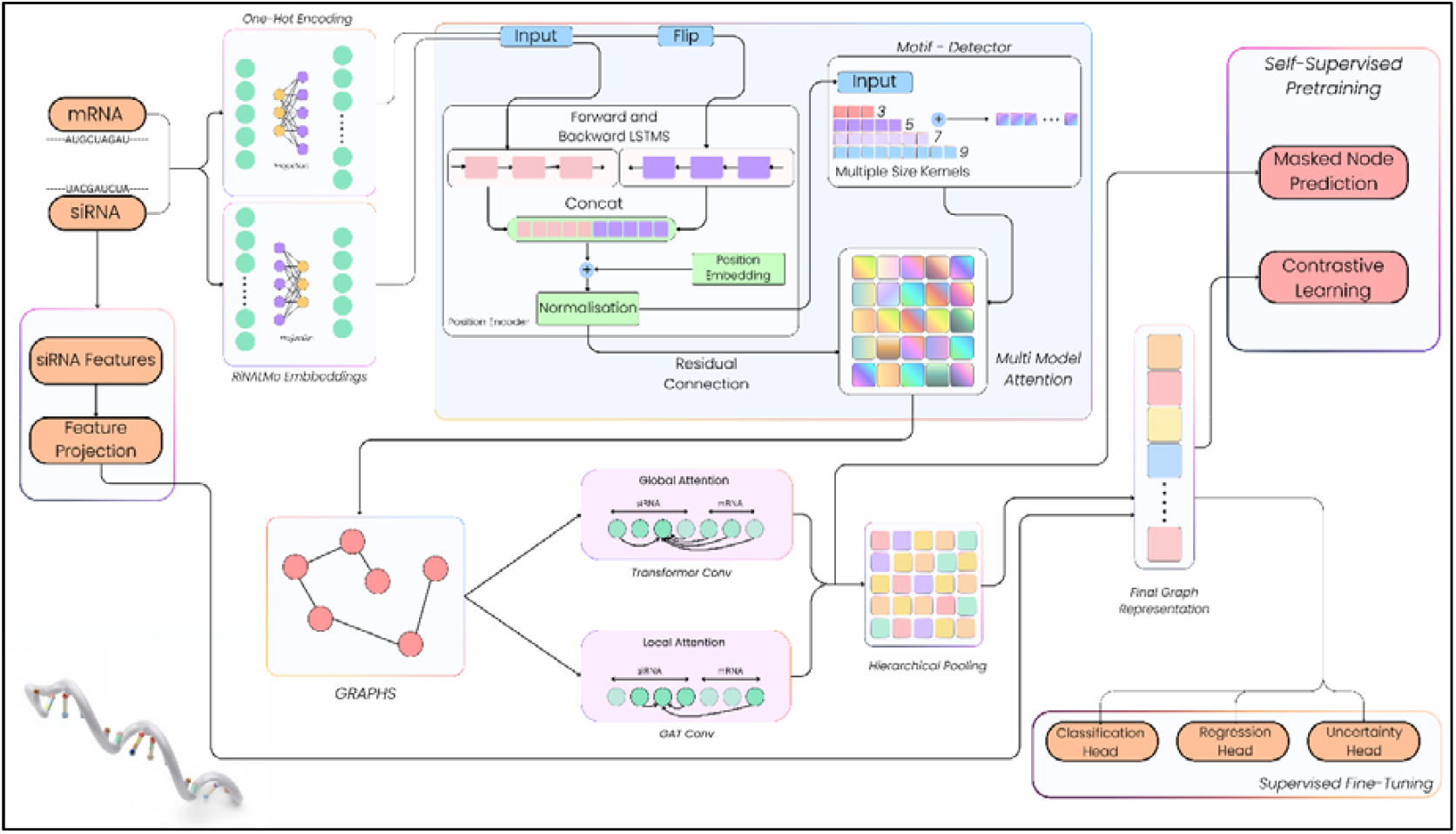
Overview of OligoGraph model architecture. siRNA and mRNA sequences are encoded using one-hot encoding, RNA language model embeddings, and engineered features, which are processed through bidirectional LSTM layers, motif detection, and attention mechanisms to capture sequence and positional dependencies. In parallel, graph representations are constructed and analyzed using transformer-based global attention, graph attention networks, and hierarchical pooling to capture structural relationships. The integrated graph representation is refined through self-supervised pretraining and used for supervised fine-tuning via classification, regression, and uncertainty heads to predict RNA efficacy.

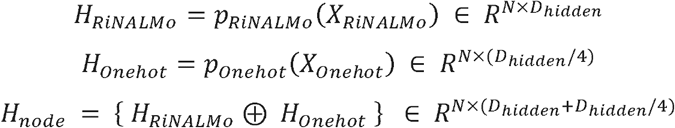

where, *X*_*RiNALMo*_ is the RiNALMo model embeddings, *X*_*Onehot*_ is the one-hot embedding, N is the number of nucleotides of both siRNA and the target site of the mRNA, *p*_*RiNALMo*_ and *p*_*Onehot*_ are dedicated projection blocks where each block consists of linear layers, layer normalization and dropout. *D*_*hidden*_ is the model’s primary hidden dimension (e.g 512 etc), *H*_*RiNALMo*_ and *H*_*Onehot*_ are the projected embeddings, *H*_*node*_ is the final node wise embedding. The operator ⊕ denotes the vector concatenation operation along the feature dimension.

#### 2.2.2. Position-aware encoder

After the input feature processing, the node-wise embeddings (*H*_*node*_) are then passed through a position-aware bidirectional encoding layer where the model utilizes two separate unidirectional LSTM networks to capture both 5’-to-3’ and 3’-to-5’ dependencies and get sequence-aware embeddings (*H*_*forward*_ and *H*_*backward*_). The output is then concatenated with node-wise learnable positional embeddings (*H*_*pos*_) and passed through layer normalization to get our position-aware embeddings (*H*_*pos-aware*_). This position-aware encoding module is critical for understanding both sequential context and positional awareness of RNA structure and function.

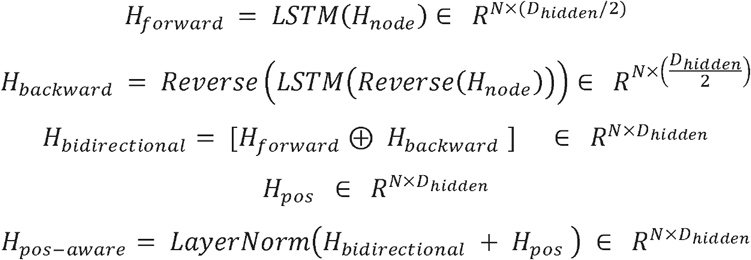

#### 2.2.3. Convolutional motif detector

The positional aware node-wise embedding (*H*_*pos-aware*_) is first transposed into 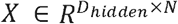 and then passed through a convolutional motif detector module with four parallel independent 1D convolutional blocks with kernel size *k* ∈ {3,5,7,9}. Each block applies 1D convolution followed by batch normalization and a ReLU activation. The output for each kernel *k, Hk* is computed as,

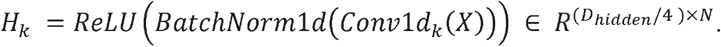

Each block has a channel dimension of 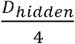 resulting in four feature maps *H*_3_,*H*_5_,*H*_7_,*H*_9_ each with 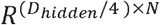 dimensions. These outputs are concatenated along the channel dimension to reassemble the full dimension feature map (*H*_*motif-pre*_).

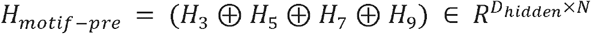

The tensor is transposed back to sequence first format to get our final output, 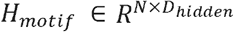

This module is useful for extracting short motifs to longer structural patterns across the RNA sequence.

#### 2.2.4. Multi-modal attention

The multi-model attention module uses a learnable attention mechanism to mitigate information loss and effectively fuse position-encoded embeddings and motif-detected embeddings that act as a learnable residual fusion layer; this preserves the sequence-aware embeddings via a skip-connection which is fused with the local structural features extracted by the convolutional motif detector. Unlike static concatenation, this mechanism uses a scalar attention mechanism to dynamically weigh the contribution of raw sequence context and processed structural motifs. The representations are projected into a latent space and a weighted residual sum is computed to get our fused embedding.

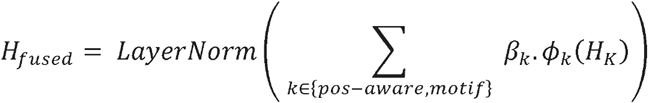

Where *ϕ*(·) is a modality-specific linear projection *Wx* + *b*. The attention coefficients *β* are derived through a Softmax operation over a learnable parameter vector *α* ∈ *R*^2^:

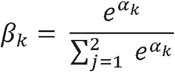

This module effectively serves as a weighted residual block, retaining sequential information for accurate predictions.

#### 2.2.5. Graph convolutional layers

To capture RNA structural dependencies, we process the fused embeddings (*H*_*fused*_) through Hybrid Graph Convolutional layers. This module utilizes complementary inductive biases by using two distinct message-passing mechanisms in parallel: Graph Transformer Convolution (TransformerConv) and Graph Attention Networks (GATConv). The TransformerConv integrates edge features to model interaction semantics, and GATConv focuses on the anisotropic aggregation of local neighborhood information.

##### 2.2.5.1. TransformerConv

Graph Transformer Convolution (TransformerConv) is a graph-native adaptation of the Transformer architecture that is used to capture high-order structural and long-range interactions within the siRNA-mRNA duplex. Unlike standard isotropic graph convolutions, TransformerConv utilizes a multi-head attention mechanism that integrates the 14-dimensional edge attribute vector *e*_*i,j*_ representing thermodynamic stability, pairing types (canonical/wobble), and backbone continuity into the attention score computation. For a given attention head *k*, the attention coefficient 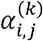 between a central nodes *i* and a neighboring node *j* ∈ *N*(*i*) is calculated as:

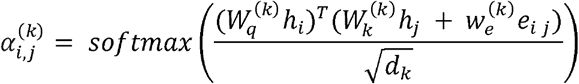

Where *h*_*i*_,*h*_*j*_, ∈ *R*^*d*^ denote the node features, 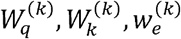 are learnable projection matrices for queries, keys, and edge attributes, respectively.

This ensures that the attention weights are dynamically modulated by the physicochemical properties of the edges. The node representations are updated through a weighted aggregation of neighbor values and edge features, followed by a gated residual connection to regulate the information flow.

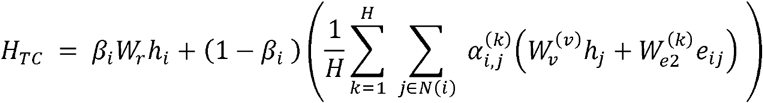

Here, *βi* is a learnable gating coefficient; H is the number of attention heads and *Wr* ensures dimensional alignment. This gating mechanism enables the model to adaptively balance local neighborhood aggregation with the preservation of current node states, mitigating the over-smoothing problem in deep GNNs.

##### 2.2.5.2. GATConv

Graph Attention Convolution (GATConv) operates in parallel with the Graph Transformer Convolution to exploit the complementary inductive bias. While TransformerConv uses dot-product attention to model global semantic relationships, GATConv uses an additive attention mechanism that focuses on anisotropic aggregation of the local structural neighborhood. The 14-dimentional edge features *e*_*i,j*_ are concatenated with node features during attention computation. The attention coefficients 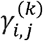 for head *k* are derived as,

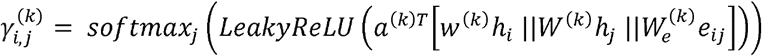

Where *a*^(*k*)^is a learnable attention vector, *W*^(*k*)^is the node projection matrix, and || denotes concatenation. This additive mechanism is effective for filtering noisy neighbors. The final node update for the GATConv is the concatenation of the multi-head outputs.

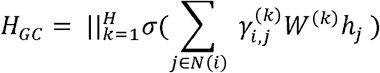

###### Hybrid inductive bias fusion

To leverage the strengths of both global semantic reasoning (TransformerConv) and local structural filtering (GATConv), we compute the node representation as the linear combination of both convolutional outputs. The final node representation (*H*_*final*_) adds the original input (*H*_*fused*_) to the computed node representation via a residual connection.

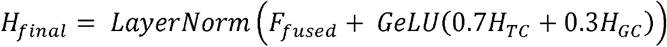

#### 2.2.6. Hierarchical pooling and siRNA feature integration

To transform the embedding from nucleotide-level representation to global duplex-level representation, a hierarchical query-based pooling mechanism was implemented. Conventional pooling aggregates all node features uniformly, diluting signals from critical functional regions such as the seed sequence. The model mitigated this by using a query-driven multi-head attention mechanism to refine node features prior to aggregation.

For graph-level aggregation, an attention-based pooling with a learned query vector 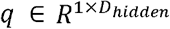 was used to attend to all node embeddings simultaneously. For the node representations 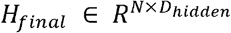, multi-head attention computes:

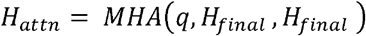

Where each head i computes scaled dot product attention:

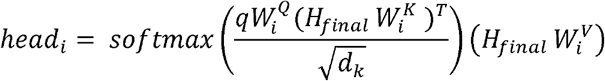

Graph-level representation via permutation-invariant pooling:

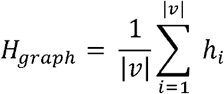

Where |v| is 38, *h*_*i*_ denotes the i-th row of *H*_*attn*_, corresponding to the enhanced embedding of node *i* after attention refinement.

The 30-dimensional siRNA thermodynamic features *f*_*thermo*_ are projected:

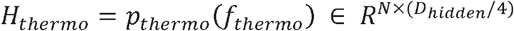

where *p*_*thermo*_ is a dedicated projection block containing linear layers.

The final graph representation is obtained by concatenating learned structural embeddings with physicochemical constraints:

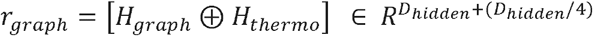

#### 2.2.7. Multi-task prediction head

The multi-modal prediction module represents the final module of the OligoGraph model. This module has two prediction heads, one for classification and one for regression.

##### Classification head

The prediction head transforms the graph representation into binary efficacy predictions using a Multi-Layer Perceptron (MLP) with GELU (Gaussian Error Linear Unit) activation and layer normalization, producing two-class logits. Focal loss is used on the classification head [30].

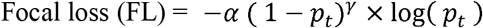

Where,

*p*_*t*_ = Model’s predicted probability for the true class.

*γ* = The focusing parameter, which is set to 2.

*α* = The balancing parameter, which is set to 1.

The goal is to effectively classify the given siRNA into two classes (ineffective and effective).

##### An Uncertainty-Aware Regression Head

The regression head takes the final graph representation and applies heteroscedastic probabilistic regression. This head consists of two MLPs with the final graph representation as inputs. The first linear map yields mean *μ* (*h*) and the second linear map yields the logarithm of the variance (*log σ*^2^ (*h*) Exponentiation ensures the variance is positive. The model treats the prediction as the mean of a Gaussian with input-dependent variance. The training uses the negative log-likelihood of this Gaussian. The loss penalizes large squared errors and a term that penalizes the predicted variance. This gives the model an adaptive way to judge errors. If the data is noisy, the model predicts higher variance, which reduces the penalty for deviations. If the model is confident, the variance must remain small, thereby increasing the penalty for errors. This provides calibrated uncertainty, allows the regression head to model noisy data more effectively, and provides uncertainty over every prediction [31].

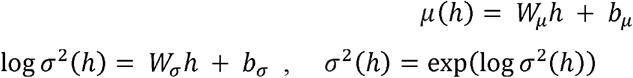

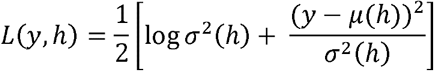

Where *h* is the final graph representation, *μ* (*h*) is the predicted mean, log *σ*^2^ (*h*)) is the predicted log variance, y is the regression label and *L*(*y,h*) is the gaussian negative log-likelihood loss.

### 2.3 Self-supervised pretraining

To address the scarcity of labeled siRNA efficacy data and to enhance the model’s capacity to learn robust molecular representations, a self-supervised pretraining framework is implemented prior to supervise fine-tuning. This phase optimizes the encoder parameters *θ* by a joint objective that combines Global Graph Contrastive Learning (GCL) and Local Masked Nucleotide Reconstruction (MNR) [32].

#### 2.3.1. Stochastic Graph Augmentation

Let *G* = (*ϑ*,*ε, X*) denote the input graph duplex, where X denotes node features. A stochastic augmentation strategy *τ* is used to generate two correlated structural views, *G*_1_, *G*_2_ ∼ *τ*(*G*) through topology and feature permutations:

- Structural corruption (Edge dropout): edges are randomly discarded from the adjacency matrix with probability *p*_*drop*_ *=* 0.15, to force the encoder to aggregate information through alternate message-passing pathways.
- Feature masking: A subset of nucleotide nodes *M* ⊂ ϑ is sampled with probability *p*_*mask*_ = 0.15. For every masked node *v* ∈ *M*, the one-hot sequence feature vector *x*_*v*_ is replaced with a zero vector 0 ∈ *R*_4_, thereby compelling the model to infer nucleotide identity from the surrounding structural context.

#### 2.3.2. Global graph contrastive learning (GCL)

For graph-level contrastive learning, a normalized temperature-scaled cross-entropy (NT-Xent) loss is used to learn the scale-invariant global representations. The encoder processes the augmented views to produce graph representations 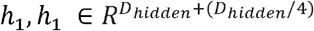 resulting from the concatenation of hierarchical pooling and projected siRNA features.

To prevent representation collapse, we map *h* to a latent metric space using a non-linear projection head 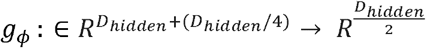:

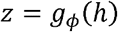

For a minibatch of *N* graphs generating 2*N* views, let *i* and *i*^+^ denote the indices of a positive pair (views from the same graph). The contrastive loss is defined as:

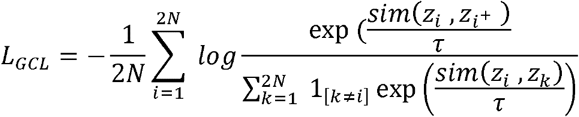

Where *τ* = 0.1 is the temperature scaling parameter, 1 is the indicator function, and 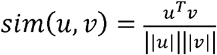 denotes cosine similarity.

#### 2.3.3. Masked Nucleotide Reconstruction (MNR)

To preserve sequence fidelity, we concurrently optimize a node-level reconstruction task. A node prediction head *q*_*φ*_ maps the embeddings 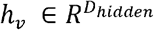 of masked nodes to a four-dimensional nucleotide probability space.

The reconstruction loss is computed as the average cross-entropy over masked nodes across both augmented views *j* ∈ {1,2}:

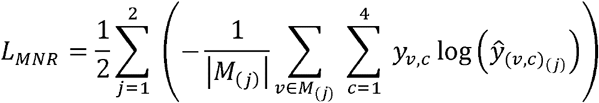

Where 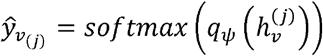 is the predicted probability distribution and *y*_*v,c*_ is the true binary label for nucleotide class c.

#### 2.3.4. Joint optimization

The final pretraining loss is the weighted summation of GCL and MNR losses:

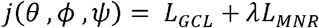

The balancing coefficient *λ* was set to 0.5. The model is optimized using AdamW with a cosine annealing learning rate scheduler and gradient clipping to ensure numerical stability.

## 3. Results and discussion

### 3.1. Intra-dataset validation

The OligoGraph (19-nucleotide version) model is compared with other siRNA efficacy predicting models like Oligoformer, Monopoli-RF, OligoWalk, siRNAPred, i-score, s-Biopredsi, DSIR etc. The metrics employed for measuring model performance were AUC (area under curve), PRC (area under the precision-recall curve), F1 score, and PCC (Pearson correlation coefficient) and given in the Table 2.

**Table 2.** Intra dataset validation.

| Methods | Huesken |  |  |  | Mixset |  |  |  | Takayuki |  |  |  |
| --- | --- | --- | --- | --- | --- | --- | --- | --- | --- | --- | --- | --- |
|  | AUC | PRC | F1 | PCC | AUC | PRC | F1 | PCC | AUC | PRC | F1 | PCC |
| **Monopol**<br>**i-RF** | 0.805 | 0.789 | 0.727 | 0.573 | 0.798 | 0.762 | 0.700 | 0.563 | 0.775 | 0.649 | 0.090 | 0.557 |
| **OligoW**<br>**alk** | 0.803 | 0.795 | 0.708 | 0.584 | 0.791 | 0.779 | 0.736 | 0.619 | 0.797 | 0.576 | 0.454 | 0.582 |
| **siRNAP**<br>**red** | 0.668 | 0.658 | 0.609 | 0.380 | 0.601 | 0.625 | 0.537 | 0.417 | 0.509 | 0.259 | 0.266 | 0.090 |
| **iScore** | 0.831 | 0.781 | 0.189 | 0.631 | 0.746 | 0.746 | 0.219 | 0.558 | 0.769 | 0.544 | 0.075 | 0.551 |
| **s-**<br>**Biopreds**<br>**i** | 0.797 | 0.773 | 0.629 | 0.662 | 0.737 | 0.758 | 0.593 | 0.520 | 0.757 | 0.535 | 0.437 | 0.528 |
| **DSIR** | 0.847 | 0.757 | 0.629 | 0.684 | 0.749 | 0.757 | 0.690 | 0.586 | 0.770 | 0.546 | 0.542 | 0.581 |
| **OligoFor**<br>**mer** | 0.861 | 0.809 | 0.758 | 0.711 | 0.845 | 0.885 | 0.764 | 0.656 | 0.862 | 0.758 | 0.576 | 0.659 |
| **OligoGr**<br>**aph** | **0.922** | **0.922** | **0.837** | **0.794** | **0.899** | **0.905** | **0.826** | **0.801** | **0.958** | **0.927** | **0.923** | **0.897** |

Of the metrics displayed in Table 2, only the OligoGraph and OligoFormer models were tested in-house; the metrics for the remaining were taken from the OligoFormer’s research paper, as the metrics we obtained were identical. The models here were trained on Huesken, Mixset, and Takayuki datasets and tested on Huesken, Mixset, and Takayuki datasets.

OligoGraph demonstrates superior metrics in the intra-dataset validation, achieving an AUC of 0.9224 and PCC of 0.794 on the Huesken dataset significantly surpassing the current best transformer-based OligoFormer (AUC 0.8619, PCC 0.7114) and the regression based DSIR (AUC 0.8472, PCC 0.6846). These results confirm that modelling the siRNA-mRNA interactions as a graph, capturing intra-strand backbones and inter-strand Watson-Crick pairings yield a higher fidelity representation than linear sequence modelling.

### 3.2. Inter-dataset validation

OligoGraph is validated on the unseen data on both the 19-nucleotide and 21-nucleotide version.

#### 3.2.1. 19-nucleotide siRNA dataset validation

In addition to intra-dataset validation, we have also tested our model’s inter-dataset performance. As the model’s performance on unseen datasets is even more crucial for evaluating robustness and generalization. We have tested the model on unseen data such as Mixset and Takayuki datasets and the metrics are presented in Table 3 and 4 and shown in Figure 2.

**Table 3.** Mixset inter-dataset validation.

| Methods | Mixset |  |  |  |
| --- | --- | --- | --- | --- |
|  | AUC | PRC | F1 | PCC |
| **Monopoli-RF** | 0.7439 | 0.7388 | 0.7002 | 0.4628 |
| **OligoWalk** | 0.7081 | 0.6998 | 0.6258 | 0.5592 |
| **siRNAPred** | 0.5782 | 0.618 | 0.5245 | 0.2062 |
| **iScore** | 0.7462 | 0.7565 | 0.2197 | 0.3687 |
| **s-Biopredsi** | 0.7374 | 0.7557 | 0.5936 | 0.3849 |
| **DSIR** | 0.7483 | 0.7637 | 0.6945 | 0.503 |
| **OligoFormer** | 0.8167 | 0.8149 | 0.7693 | 0.5879 |
| **OligoGraph** | **0.8257** | **0.8260** | 0.7481 | **0.6150** |

**Table 4.** Takayuki inter-dataset validation.

| Methods | Takayuki |  |  |  |
| --- | --- | --- | --- | --- |
|  | AUC | PRC | F1 | PCC |
| **OligoFormer** | 0.5845 | 0.3488 | 0.3803 | 0.2008 |
| **OligoGraph** | **0.6960** | **0.4216** | **0.6055** | **0.4566** |

**Figure 2.**
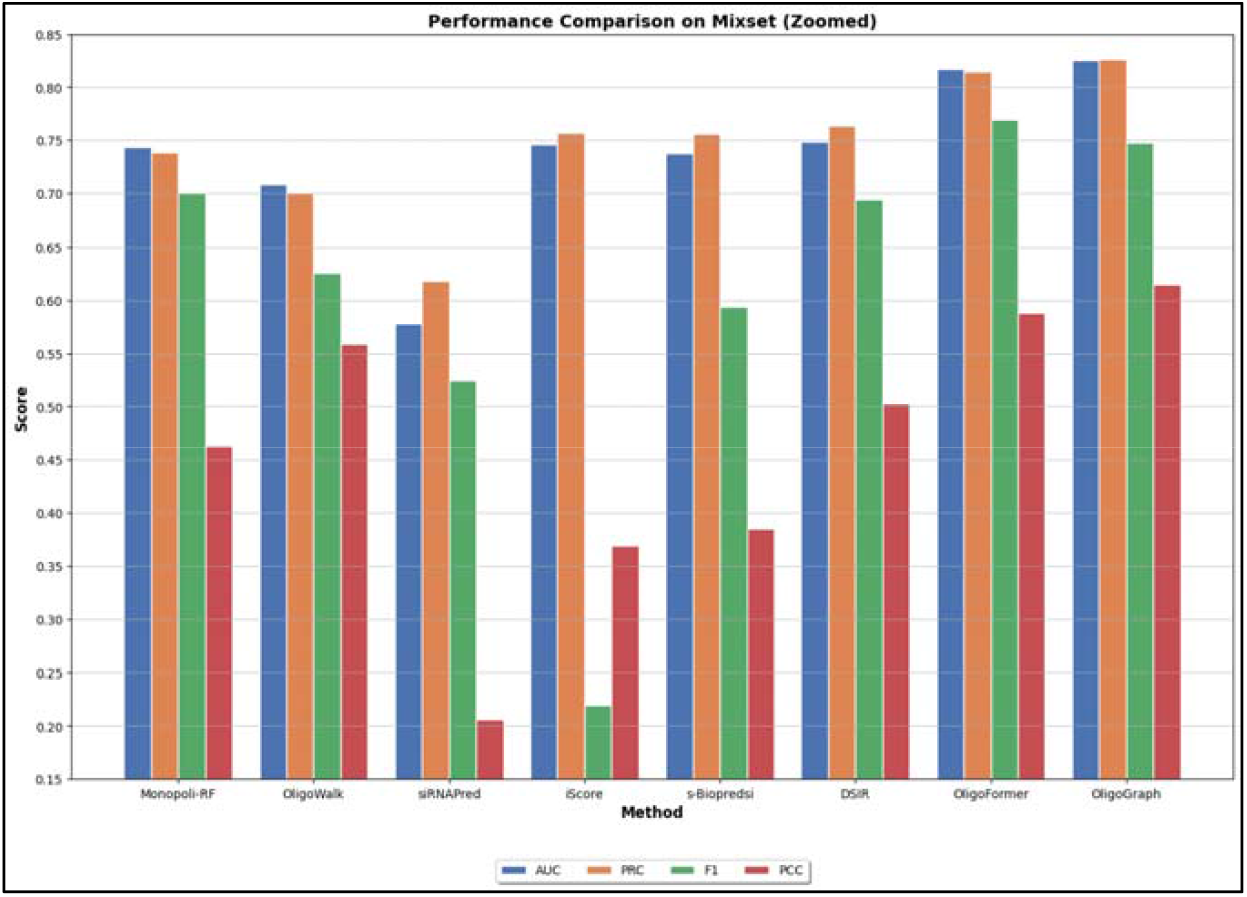
Performance comparison of machine learning and deep learning models on the MixSet dataset. The figure 2 compares the predictive performance of different siRNA efficacy prediction methods, including Monopoli-RF, OligoWalk, siRNAPred, iScore, s-Biopredsi, DSIR, OligoFormer, and OligoGraph, evaluated on the MixSet dataset. Performance is measured using four metrics: Area Under the ROC Curve (AUC), Precision-Recall Curve (PRC), F1 score, and Pearson Correlation Coefficient (PCC). OligoGraph demonstrates superior overall performance across all evaluation metrics, indicating its improved ability for accurate siRNA efficacy prediction compared to existing approaches.

Of the metrics displayed above in Table 3, only the OligoGraph and OligoFormer models were tested in-house, and the metrics for the rest were taken from the OligoFormer’s research paper as the metrics we obtained for OligoFormer were identical. The metrics displayed in bold are the best metrics obtained after multiple training runs. The models were trained on the Huesken dataset and validated on the mixset.

The models were trained on the Huesken dataset and validated on the Takayuki dataset are presented in Table 4. No metrics for prior models to OligoFormer were displayed as they weren’t tested on Takayuki dataset.

Inter-dataset benchmarking on 19-nucleotide sequences confirms the model’s generalization and robustness against data distribution shifts. On the unseen Mixset, OligoGraph maintains a PCC of 0.6150 outperforming OligoFormer (PCC 0.5879). In the challenging Takayuki dataset OligoGraph (AUC 0.6654, PCC 0.3959) outperformed OligoFormer (AUC 0.5845, PCC 0.3959). This difference suggests that the model does not overfit to specific experimental artifacts and has superiour generalization compared to existing models.

#### 3.2.2. 21-nucleotide siRNA dataset validation

The model here was trained on the 21-nucleotide HUVK dataset and tested on the 21-nucleotide Simone dataset. The metrics are presented in Table 5.

**Table 5.** Simone dataset.

| Methods | Simone |  |
| --- | --- | --- |
|  | AUC | PCC |
| **i-Score** | 0.623 | 0.379 |
| **s-Biopredsi** | 0.628 | 0.372 |
| **DSIR** | 0.649 | 0.386 |
| **CNN** | 0.660 | 0.376 |
| **GNN4siRNA** | 0.612 | 0.262 |
| **siRNA discovery** | 0.702 | 0.464 |
| **OligoGraph** | 0.7204 | 0.4949 |

Table 5 presents the metrics of siRNA discovery and prior models as per the siRNA discovery research paper. The models were trained on the HUVK dataset and tested on the Simone dataset. OligoGraph obtained 0.7204 AUC-ROC, 0.8551 AUC-PRC, 0.6892 F1 score, 0.4949 PCC, and an MSE of 0.0421 on the unseen Simone dataset. These metrics establish the OligoGraph’s generalization and robustness on unseen data.

### 3.3. Ablation study

To precisely assess the importance and contribution of each key component in the OligoGraph architecture, we conducted a comprehensive set of ablation studies to measure inter-dataset performance. The metrics are presented in the Table 6 and shown in Figure 3.

**Table 6.** Ablation study for different modules.

| **Ablation** | **F1 score** | **AUC ROC** | **AUC PR** | **PCC** |
| --- | --- | --- | --- | --- |
| **No GAT Conv** | 0.7282 | 0.8023 | 0.8180 | 0.5799 |
| **No RiNALMo** | 0.7435 | 0.8118 | 0.8249 | 0.5860 |
| **No Trans Conv** | 0.6773 | 0.7250 | 0.7350 | 0.3956 |
| **No siRNA Features** | 0.6815 | 0.8093 | 0.8296 | 0.5835 |
| **No Bilstm** | 0.7466 | 0.8126 | 0.8320 | 0.5918 |
| **Separate Processing** | 0.7240 | 0.7979 | 0.8020 | 0.5683 |
| **Complete OligoGraph** | 0.7481 | 0.8257 | 0.8260 | 0.6150 |

**Figure 3.**
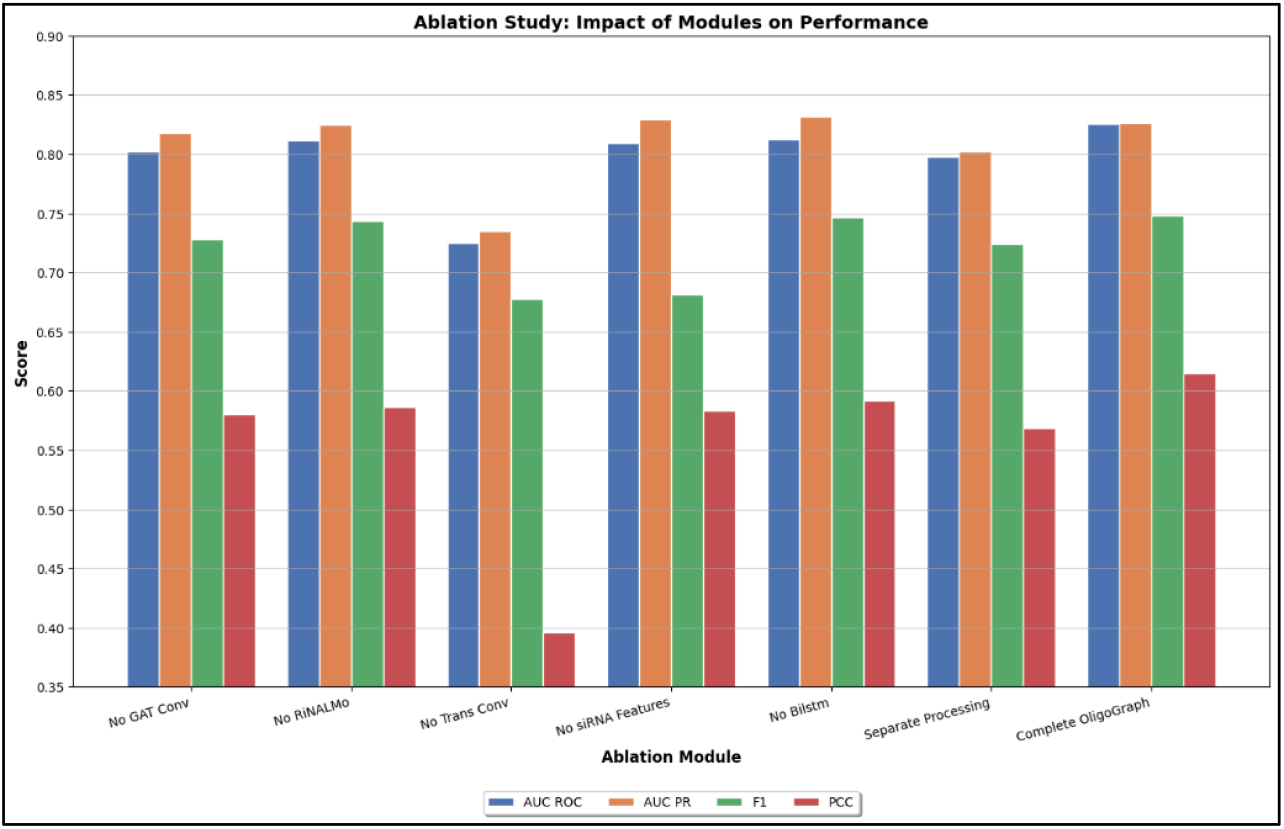
Ablation study showing the impact of individual modules on OligoGraph performance. The figure 3 presents an ablation analysis evaluating the contribution of key modules, including GAT convolution, RiNALMo, transformer convolution, engineered siRNA features, and BiLSTM layers, to the overall performance of OligoGraph. Model performance is assessed using AUC-ROC, AUC-PR, F1 score, and Pearson Correlation Coefficient (PCC) after removing or modifying each component. The complete OligoGraph model achieves the highest performance across all metrics, demonstrating the importance of integrating sequence embeddings, graph attention, and feature fusion for accurate siRNA efficacy prediction.

The model was trained on Huesken, tested on Mixset and the metrics are presented in Table 6.

The results clearly underscore the necessity of the proposed multi modal and structural feature integration approach. The most significant performance drop was observed when the Transformer Convolutional Layer was removed, resulting in a large decrease in the Pearson Correlation Coefficient (PCC) from 0.6150 to 0.3956 and the AUC-ROC to 0.7250. This highlights the fundamental role of graph-tuned Scaled dot product attention mechanism in effectively gathering complete sequence information, which is critical for accurate efficacy prediction.

The removal of the siRNA Features led to a significant drop in F1 score from 0.7481 to 0.6815 and AUC-PR, confirming that explicit siRNA features are not merely redundant but necessary for reliable binding affinity modeling. Similarly, ablating the GAT convolutional Layer led to a notable reduction across all the metrics, with AUC-ROC dropping to 0.8023. This validates our design choice of using a weighted combination of TransformerConv and GATConv, where the GAT layer provides complementary structural neighborhood awareness that the pure self-attention mechanism might overlook.

The approach of separate processing this ablation study validated the superiority of the preprocessing approach used in the complete model by implementing a separate processing strategy as a comparison. Models were pretrained on Huesken and Takayuki, trained on Huesken and validated on Mix. The separate processing approach fed each 19 nucleotide strand into RiNALMo independently, generated separate embeddings then concatenate them with an additional strand type indicator (0 for siRNA and 1 for mRNA), achieving 0.7979 AUC and confirming that isolation strands degrades performance. While the concatenated approach used in the complete model combined siRNA and mRNA into a single 38 token input before passing through RiNALMo achieving 0.8257 AUC. This gap shows that RiNALMo’s transformer architecture captures critical inter strand communication when processing the combined sequences with attention mechanisms encoding the cross strand interactions. The separate processing method crashed the F1 score to 0.7240 and Pearson Correlation to 0.5683 validating that leveraging RiNALMo’s capacity for inter-molecular dependencies is essential for accurate siRNA efficacy prediction.

The removal of Position Encoded BiLSTM lead to a minor drop of around 1 percent showing that they do provide subtle information which improves the accuracy. The removal of RiNALMo pre-trained features and replacing it with learnable nn.embeddings lead to a drop of around 3 percent in PCC showing that RiNALMo embeddings provides information which capture the inter stand relationships, sequence features and siRNA efficacy.

## 4. Conclusion

In this study, we presented OligoGraph, a geometric deep learning framework that advances the predictive modeling of siRNA efficacy by integrating structural, thermodynamic, and semantic features within a graph-based architecture. Unlike preceding models that rely primarily on sequence linearity or limited hand-crafted features, OligoGraph explicitly models the siRNA-mRNA duplex as a graph, capturing complex non-linear interactions through a hybrid message-passing mechanism.

The synergistic application of TransformerConv and GATConv layers proved critical for capturing both long-range dependencies and local structural neighborhoods, as evidenced by the substantial performance degradation observed in their absence during ablation studies. Furthermore, incorporating RiNALMo foundation model embeddings and specific thermodynamic constraints enabled the model to overcome the limitations of data scarcity and the inherent bias in publicly available datasets.

Model validation across intra-dataset and inter-dataset benchmarks demonstrates that OligoGraph achieves commendable performance, significantly outperforming existing fourth-generation tools such as OligoFormer and legacy algorithms like DSIR. The model exhibits superior generalization, particularly in the challenging transfer-learning scenarios represented by the Takayuki and Mixset datasets. By effectively fusing self-supervised pretraining with uncertainty-aware regression and multi-modal attention, OligoGraph provides a robust, high-fidelity deep learning framework for siRNA efficacy prediction, potentially accelerating the development of effective RNAi-based therapeutics.

OligoGraph presents several promising avenues for future enhancement and expansion. Architectural improvements could include integrating more advanced messaging-passing mechanisms in graphs to better capture long-range features. Enhanced feature engineering opportunities include incorporating dynamic structural features like predicted RNA secondary structures, and 3D spatial arrangements beyond linear sequence representations. The model could be extended to predict off-target effects.

The current positional BiLSTM encoder and convolutional motif detector, while functional, provide limited added value and represent opportunities for improvement through more advanced sequence modeling approaches such as temporal convolutional networks or transformer-based positional encoders that could better capture long-range dependencies and complex motif patterns.

Additionally, exploration of attention mechanisms specifically designed for biological sequence data that incorporate evolutionary conservation patterns could enhance model performance.

## Abbreviations

AGO2: Argonaute-2
AUC: Area Under Curve
BiLSTM: Bidirectional Long Short Term Memory
CNN: Convolutional Neural Network
dsRNA: double-stranded Ribonucleic Acid
GATConv: Graph Attention Convolution
GNN: Graph Neural Network
MLP: Multilayer Perceptron
mRNA: Messenger Ribonucleic Acid
MSE: mean Squared Error
nt: Nucleotide
nn: Neural Network
PCC: Pearson Correlation Coefficient
PR: Precision-Recall
RISC: RNA-Induced Silencing Complex
RNA: Ribonucleic Acid
RNAi: Ribonucleic Acid interference
RNase: Ribonuclease
ROC: Receiver Operating Characteristic
siRNA: small-interfering Ribonucleic acid
TransConv: Transformer Convolution

## DECLARATIONS

### Ethics approval and consent to participate

Not applicable

### Consent for publication

Not applicable

### Data availability

All data used and analyzed in this study are provided within the manuscript.

### Code availability

The code is available at GitHub Link: https://github.com/drugparadigm/OligoGraph

### Author Contributions

**S.S.S. and V.V.K**.: Investigation; validation; methodology; visualization; writing-original draft; formal analysis. **S.R.S**.: Data curation; formal analysis; methodology; validation; visualization. **B.B.C**.: Conceptualization; methodology; validation; visualization; formal analysis; data curation; software; supervision; formal analysis; writing-review and editing. **V.K**.: Conceptualization; methodology; writing-review and editing; supervision; formal analysis.

## Acknowledgements

We, the authors, Sriram Shravan Saligram, Vishnu Vardhan Kasturi, Sidhardha Reddy Surkanti, Basangari Bhargava Chary, and Vani Kondaparthi, express our sincere gratitude to the Drugparadigm Research Lab for providing the necessary facilities and infrastructure that enabled the successful completion of this work.

## REFERENCES

1. Khvorova A, Reynolds A, Jayasena SD. Functional siRNAs and miRNAs exhibit strand bias. Cell. [Comparative Study]. 2003;115(2):209–16. DoI:10.1016/s0092-8674(03)00801-8

2. Elbashir SM, Harborth J, Lendeckel W, Yalcin A, Weber K, Tuschl T. Duplexes of 21-nucleotide RNAs mediate RNA interference in cultured mammalian cells. Nature. [Research Support, Non-U.S. Gov’t]. 2001;411(6836):494–8. DoI:10.1038/35078107

3. Liu J, Carmell MA, Rivas FV, Marsden CG, Thomson JM, Song JJ, et al. Argonaute2 is the catalytic engine of mammalian RNAi. Science. [Research Support, Non-U.S. Gov’t Research Support, U.S. Gov’t, P.H.S.]. 2004;305(5689):1437–41. DoI:10.1126/science.1102513

4. Sini P, Cannas A, Koleske AJ, Di Fiore PP, Scita G. Abl-dependent tyrosine phosphorylation of Sos-1 mediates growth-factor-induced Rac activation. Nat Cell Biol. [Research Support, Non-U.S. Gov’t]. 2004;6(3):268–74. DoI:10.1038/ncb1096

5. Reynolds A, Leake D, Boese Q, Scaringe S, Marshall WS, Khvorova A. Rational siRNA design for RNA interference. Nat Biotechnol. [Comparative Study Evaluation Study Research Support, U.S. Gov’t, Non-P.H.S.]. 2004;22(3):326–30. DoI:10.1038/nbt936

6. Ui-Tei K, Naito Y, Takahashi F, Haraguchi T, Ohki-Hamazaki H, Juni A, et al. Guidelines for the selection of highly effective siRNA sequences for mammalian and chick RNA interference. Nucleic Acids Res. [Guideline Research Support, Non-U.S. Gov’t]. 2004;32(3):936–48. DoI:10.1093/nar/gkh247

7. Amarzguioui M, Prydz H. An algorithm for selection of functional siRNA sequences. Biochem Biophys Res Commun. [Comparative Study Evaluation Study Research Support, Non-U.S. Gov’t Validation Study]. 2004;316(4):1050–8. DoI:10.1016/j.bbrc.2004.02.157

8. Huesken D, Lange J, Mickanin C, Weiler J, Asselbergs F, Warner J, et al. Design of a genome-wide siRNA library using an artificial neural network. Nat Biotechnol. [Evaluation Study]. 2005;23(8):995–1001. DoI:10.1038/nbt1118

9. Ichihara M, Murakumo Y, Masuda A, Matsuura T, Asai N, Jijiwa M, et al. Thermodynamic instability of siRNA duplex is a prerequisite for dependable prediction of siRNA activities. Nucleic Acids Res. [Research Support, Non-U.S. Gov’t Validation Study]. 2007;35(18):e123. DoI:10.1093/nar/gkm699

10. Lu ZJ, Mathews DH. OligoWalk: an online siRNA design tool utilizing hybridization thermodynamics. Nucleic Acids Res. [Research Support, N.I.H., Extramural]. 2008;36(Web Server issue):W104–8. DoI:10.1093/nar/gkn250

11. Filhol O, Ciais D, Lajaunie C, Charbonnier P, Foveau N, Vert JP, et al. DSIR: assessing the design of highly potent siRNA by testing a set of cancer-relevant target genes. PLoS One. [Research Support, Non-U.S. Gov’t]. 2012;7(10):e48057. DoI:10.1371/journal.pone.0048057

12. Qureshi A, Thakur N, Kumar M. VIRsiRNApred: a web server for predicting inhibition efficacy of siRNAs targeting human viruses. J Transl Med. [Research Support, Non-U.S. Gov’t Validation Study]. 2013;11:305. DoI:10.1186/1479-5876-11-305

13. Dar SA, Gupta AK, Thakur A, Kumar M. SMEpred workbench: A web server for predicting efficacy of chemicallymodified siRNAs. RNA Biol. 2016;13(11):1144–51. DoI:10.1080/15476286.2016.1229733

14. Cha B, Geng X, Mahamud MR, Zhang JY, Chen L, Kim W, et al. Complementary Wnt Sources Regulate Lymphatic Vascular Development via PROX1-Dependent Wnt/beta-Catenin Signaling. Cell Rep. [Research Support, N.I.H., Extramural Research Support, Non-U.S. Gov’t]. 2018;25(3):571–84 e5. DoI:10.1016/j.celrep.2018.09.049

15. La Rosa M, Fiannaca A, La Paglia L, Urso A. A Graph Neural Network Approach for the Analysis of siRNA-Target Biological Networks. Int J Mol Sci. 2022;23(22). DoI:10.3390/ijms232214211

16. Long R, Guo Z, Han D, Liu B, Yuan X, Chen G, et al. siRNADiscovery: a graph neural network for siRNA efficacy prediction via deep RNA sequence analysis. Brief Bioinform. 2024;25(6). DoI:10.1093/bib/bbae563

17. Bai Y, Zhong H, Wang T, Lu ZJ. OligoFormer: an accurate and robust prediction method for siRNA design. Bioinformatics. 2024;40(10). DoI:10.1093/bioinformatics/btae577

18. Fonck C, Su C, Arens J, Koziol E, Srimani J, Henshaw J, et al. Lack of germline transmission in male mice following a single intravenous administration of AAV5-hFVIII-SQ gene therapy. Gene Ther. [Research Support, Non-U.S. Gov’t]. 2023;30(7-8):581–6. DoI:10.1038/s41434-022-00318-5

19. Penic RJ, Vlasic T, Huber RG, Wan Y, Sikic M. RiNALMo: general-purpose RNA language models can generalize well on structure prediction tasks. Nat Commun. 2025;16(1):5671. DoI:10.1038/s41467-025-60872-5

20. Katoh T, Suzuki T. Specific residues at every third position of siRNA shape its efficient RNAi activity. Nucleic Acids Res. [Research Support, Non-U.S. Gov’t]. 2007;35(4):e27. DoI:10.1093/nar/gkl1120

21. Amarzguioui M, Holen T, Babaie E, Prydz H. Tolerance for mutations and chemical modifications in a siRNA. Nucleic Acids Res. [Research Support, Non-U.S. Gov’t]. 2003;31(2):589–95. DoI:10.1093/nar/gkg147

22. Harborth J, Elbashir SM, Vandenburgh K, Manninga H, Scaringe SA, Weber K, et al. Sequence, chemical, and structural variation of small interfering RNAs and short hairpin RNAs and the effect on mammalian gene silencing. Antisense Nucleic Acid Drug Dev. [Research Support, Non-U.S. Gov’t]. 2003;13(2):83–105. DoI:10.1089/108729003321629638

23. Hsieh AC, Bo R, Manola J, Vazquez F, Bare O, Khvorova A, et al. A library of siRNA duplexes targeting the phosphoinositide 3-kinase pathway: determinants of gene silencing for use in cell-based screens. Nucleic Acids Res. [Research Support, Non-U.S. Gov’t Research Support, U.S. Gov’t, P.H.S.]. 2004;32(3):893–901. DoI:10.1093/nar/gkh238

24. Vickers TA, Koo S, Bennett CF, Crooke ST, Dean NM, Baker BF. Efficient reduction of target RNAs by small interfering RNA and RNase H-dependent antisense agents. A comparative analysis. J Biol Chem. 2003;278(9):7108–18. DoI:10.1074/jbc.M210326200

25. Sciabola S, Cao Q, Orozco M, Faustino I, Stanton RV. Improved nucleic acid descriptors for siRNA efficacy prediction. Nucleic Acids Res. [Evaluation Study Research Support, Non-U.S. Gov’t]. 2013;41(3):1383–94. DoI:10.1093/nar/gks1191

26. Xia T, SantaLucia J, Jr., Burkard ME, Kierzek R, Schroeder SJ, Jiao X, et al. Thermodynamic parameters for an expanded nearest-neighbor model for formation of RNA duplexes with Watson-Crick base pairs. Biochemistry. [Comparative Study Research Support, U.S. Gov’t, P.H.S.]. 1998;37(42):14719–35. DoI:10.1021/bi9809425

27. SantaLucia J, Jr. A unified view of polymer, dumbbell, and oligonucleotide DNA nearest-neighbor thermodynamics. Proc Natl Acad Sci U S A. [Research Support, Non-U.S. Gov’t]. 1998;95(4):1460–5. DoI:10.1073/pnas.95.4.1460

28. Turner DH, Sugimoto N, Freier SM. RNA structure prediction. Annu Rev Biophys Biophys Chem. [Research Support, U.S. Gov’t, P.H.S. Review]. 1988;17:167–92. DoI:10.1146/annurev.bb.17.060188.001123

29. Starnes SK, Horne WS, Del Valle JR. Impact of Strand Edge N-Amination on the Stability of a Parallel beta-Hairpin Fold. J Org Chem. 2025;90(48):17214–20. DoI:10.1021/acs.joc.5c02479

30. Lin T-Y, Goyal P, Girshick RB, He K, Dollár P. Focal Loss for Dense Object Detection. 2017 IEEE International Conference on Computer Vision (ICCV). 2017:2999–3007

31. Kendall A, Gal Y. What Uncertainties Do We Need in Bayesian Deep Learning for Computer Vision? ArXiv. 2017;abs/1703.04977

32. Hu W, Liu B, Gomes J, Zitnik M, Liang P, Pande VS, et al. Strategies for Pre-training Graph Neural Networks. arXiv: Learning. 2019;

